# An efficient polymer cocktail-based transportation method for cartilage tissue, yielding chondrocytes with enhanced hyaline cartilage expression during in vitro culturing

**DOI:** 10.1101/2021.07.14.452414

**Authors:** Shojiro Katoh, Hiroshi Yoshioka, Shoji Suzuki, Hiroyuki Nakajima, Masaru Iwasaki, Rajappa Senthilkumar, Senthilkumar Preethy, Samuel JK Abraham

**Affiliations:** Edogawa Evolutionary Lab of Science, Edogawa Hospital Campus, 2-24-18, Higashi Koiwa, Edogawa-Ku, Tokyo 133-0052, Japan; Department of Orthopaedic Surgery, Edogawa Hospital, 2-24-18, Higashi Koiwa, Edogawa-Ku, Tokyo 133-0052, Japan; Mebiol Inc., 1-25-8, Nakahara, Hiratsuka, 254-0075, Kanagawa, Japan; Department of Clinical Education, University of Yamanashi - Faculty of Medicine, 1110, Shimokato, Chuo, Yamanashi 409-3898, Japan; II Department of Surgery, University of Yamanashi - Faculty of Medicine, 1110, Shimokato, Chuo, Yamanashi 409-3898, Japan; Centre for Advancing Clinical Research (CACR), University of Yamanashi - Faculty of Medicine, 1110, Shimokato, Chuo, Yamanashi 409-3898, Japan; The Fujio-Eiji Academic Terrain (FEAT), Nichi-In Centre for Regenerative Medicine (NCRM), PB 1262, Chennai 600034, Tamil Nadu, India; The Mary-Yoshio Translational Hexagon (MYTH), Nichi-In Centre for Regenerative Medicine (NCRM), PB 1262, Chennai 600034, Tamil Nadu, India; JBM Inc., 3-1-14, Higashi Koiwa, Edogawa-Ku, Tokyo 133-0052, Japan; Antony-Xavier Interdisciplinary Scholastics (AXIS), GN Corporation Co. Ltd., 3-8, Wakamatsu, Kofu, Yamanashi 400-0866, Japan

**Author notes:** **Corresponding author information:** Dr. Samuel JK Abraham, Centre for Advancing Clinical Research (CACR), University of Yamanashi, Faculty of Medicine, Address for correspondence: 3-8, Wakamatsu, Kofu, 400-0866, Yamanashi, Japan, Email id-; Alternate email id.

**Keywords:** Transportation, Cartilage, Chondrocytes, Hyaline Phenotype, Thermo-reversible Gelation Polymer (TGP)

## Abstract

Chondrocytes are used in cell-based therapies such as autologous chondrocyte implantation (ACI) and matrix-associated cartilage implantation (MACI). To transport the cartilage tissue to the laboratory for in vitro culturing, phosphate-buffered saline (PBS), Euro-Collins solution (ECS) and Dulbecco’s Modified Eagle’s Medium (DMEM) are commonly employed at 4-8 °C. In this study, eight samples of human cartilage biopsy tissues from elderly patients with severe osteoarthritis undergoing arthroscopy, which would otherwise have been discarded, were used. The cartilage tissue samples were compared to assess the cell yield between two transportation groups: i) a thermo-reversible gelation polymer (TGP) based method without cool preservation (~25 °C) and ii) ECS transport at 4 °C. These samples were subjected to in vitro culture in a two-dimensional (2D) monolayer for two weeks and subsequently in a three-dimensional (3D) TGP scaffold for six weeks. The cell count obtained from the tissues transported in TGP was higher (0.2 million cells) than those transported in ECS (0.08 million cells) both after initial processing and after in vitro culturing for 2 weeks in 2D (18 million cells compared with 10 million cells). In addition, mRNA quantification demonstrated significantly higher expression of Col2a1 and SOX-9 in 3D-TGP cultured cells and lower expression of COL1a1 in RT-PCR, characteristic of the hyaline cartilage phenotype, than in 2D culture. This study confirms that the TGP cocktail is suitable for both the transport of human cartilage tissue and for in vitro culturing to yield better-quality cells for use in regenerative therapies.

## Introduction

Human primary tissue-derived cell culture is an essential part of cell-based regenerative therapies. Autologous chondrocyte implantation (ACI) and matrix-associated cartilage implantation (MACI) represent established cell-based therapies for cartilage repair [1]. Cell culturing for application in ACI and MACI requires cartilage tissue biopsy samples or surgical specimens to be transported to the laboratory from the place of collection. This, in turn, necessitates effective methodologies for preserving the viability and characteristics of the resident tissue chondrocytes for culture expansion [1]. Furthermore, transportation techniques must facilitate the transportation of clinical tissue samples from distant locations, even from overseas, while maintaining cell viability and native characteristics when samples are processed after transport for cell therapy applications [2]. Phosphate-buffered saline (PBS), Dulbecco’s Modified Eagle’s Medium (DMEM) and Euro-Collins solution (ECS) represent common benchmark tissue preservation/transportation solutions [2]. All three solutions require temperature maintenance at 4 °C during transportation and preservation. We previously reported the transportation of human corneal endothelial tissues from the collection source to the laboratory over a distance of 2500 km without the need for cool preservation by employing a thermo-reversible gelation polymer (TGP) methodology [3]. In a later study, similar corneal endothelial tissues were processed to derive human corneal endothelial precursor cells that were employed in human pilot clinical studies to treat bullous keratopathy [4]. Human retinal pigment epithelial tissue has also been transported using the TGP methodology [5]. TGP remains a gel at temperatures above sol-gel transition temperatures (~20 °C) and liquefies at lower temperatures [6]. Therefore, tissues and cells are placed in the TGP when it is in liquid form (refrigerated until use) and can then be transported without the need for cool preservation at temperatures ranging from 25 to 50 °C, with the tissues and cells preserved in the TGP gel [3]. The TGP is in a lyophilized form and can be reconstituted with any culture medium [6] specific to the tissue/cell being transported, and even saline, which is employed during clinical transplantation [7]. We previously reported the ability to tissue engineer cartilage using a three-dimensional (3D) TGP scaffold-based culture of chondrocytes from non-weight-bearing cartilage [8,9] and chondrocytes derived from elderly donors with osteoarthritis [10]. We also reported that the 3D-TGP tissue-engineered cartilage contained cells positive for pluripotency markers in lectin microarray [10]. In this study, we analysed the possibility of transporting discarded cartilage biopsy tissue obtained from elderly osteoarthritic donors during arthroscopy using the TGP-based preservation method to determine whether it can maintain viability and yield native hyaline phenotypic cartilage tissue when used for in vitro cell culturing.

## Methods

The study was approved by the institutional ethics committee of Edogawa Hospital, Tokyo, Japan. Eight samples (n = 8) of cartilage biopsy tissues weighing 0.1 to 0.3 g and derived from elderly patients (aged 60-85 years) with severe osteoarthritis undergoing arthroscopy, which would have otherwise been discarded, were employed in the study. The study was conducted in accordance with relevant guidelines and regulations, and informed consent was obtained from all participants and/or their legal guardians. The cartilage tissues obtained were divided into two equal portions by visual observation. One portion was placed in 5 ml of ECS solution (Figure 1A), whereas the other was placed in 5 ml of liquefied TGP reconstituted in DMEM in a 1:9 ratio (Figure 1B). To prepare the TGP cocktail for transportation, lyophilized TGP vials (1 g) were obtained from M/s GN Corporation, Japan (manufactured by Mebiol, Inc., Japan). One gram of TGP was reconstituted with 9 ml of DMEM and incubated at 4 °C overnight. TGP, when reconstituted, becomes a gel and remains such at temperatures above the sol-gel transition temperature of 20 °C, whereas it liquefies at temperatures below 20 °C.

**Figure 1:**
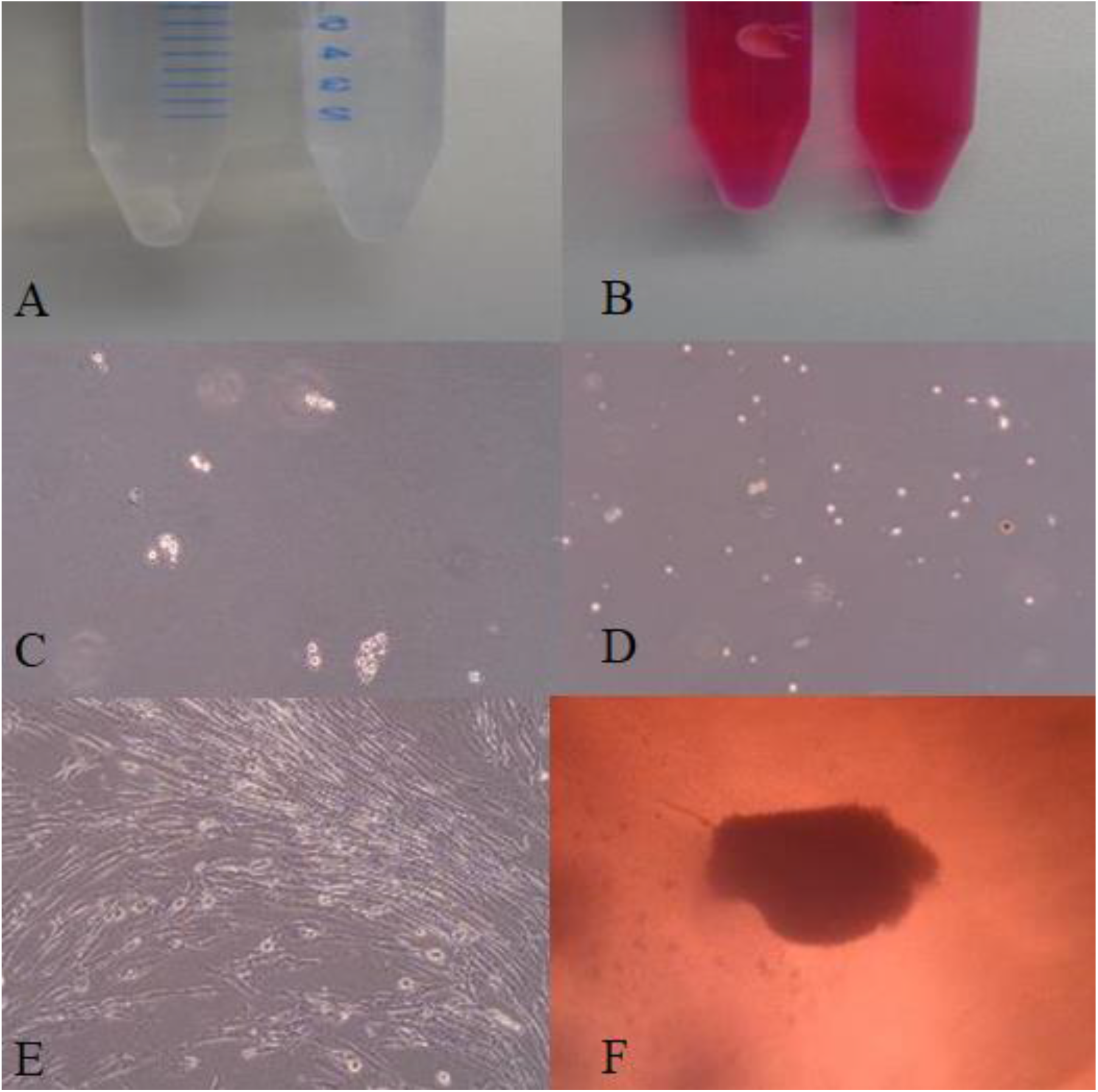
A. Cartilage biopsy tissues transported in ECS; B. Cartilage biopsy tissues transported in TGP; C. Cells derived from ECS-transported tissue after isolation; D. Cells derived from TGP-transported tissue after isolation; E. Fibroblast dedifferentiated chondrocytes during 2D-monolayer culture; F. Cartilage tissue construct in 3D-TGP culture.

The containers were closed and sealed, and the ECS container was placed in transportation boxes containing thermo packs that maintained the temperature at 4 °C, whereas the TGP container was placed in transportation boxes with thermo packs that maintained the temperature around 25 °C (to avoid freezing during the sub-zero outside temperatures that occur in Japan during winter) (Japan Meteorological Agency) [11]. The temperature and other parameters pertaining to transporting the samples in TGP are described in Table 1. The duration of transportation from the hospital where the cartilage biopsy was taken to the cell processing laboratory was 1 to 1.5 hours. Upon reaching the laboratory, the samples were subjected to chondrocyte isolation and culture following the methodology we reported earlier [8-10]. Briefly, the cartilage tissue was weighed and subjected to digestion with 0.25% trypsin for 30 min in an orbital shaker at 150 rpm at 37 °C. The tissues were then subjected to 2 mg/ml collagenase digestion for 12-18 h at 37 °C in an orbital shaker, followed by filtering with a 100-μm cell strainer. After the undigested tissues were discarded, the filtrate was centrifuged at 1000 rpm for 10 min. The cells obtained were counted using the trypan blue dye exclusion method. After two-dimensional (2D) monolayer culturing in media containing low-glucose DMEM, 10% autologous plasma, 1% penicillin streptomycin, 50 μg/ml gentamicin and 0.25 μg/ml amphotericin B and L-ascorbic acid (50 mg/ml) for 2 weeks at 37 °C with 5% CO_2_, the cells were seeded into the 3D-TGP scaffold using the same media composition and cultured for six weeks at 37 °C with 5% CO_2_ in an orbital shaker. The chondrocytes cultured in a 2D monolayer and the 3D-TGP tissue-engineered cartilage tissue was subjected to qRT-PCR for the expression of SOX9, COL1a1 and COL2a1. Total RNA was isolated using the NucleoSpin RNA XS kit (Macherey-Nagel). RT-PCR was performed using the One Step TB Green PrimeScript PLUS RT-PCR Kit (Perfect Real Time; Takara, Japan) and Thermal Cycler Dice^®^ Real Time System Lite (TP700, TaKaRa). The primers employed are provided in Table 2. For mRNA quantification, we used the delta-delta Ct approximation method, which measures the relative levels of mRNA between two samples by comparing them with a second RNA that serves as a normalization standard. GADPH was used as the normalization standard. Excel software statistics package analysis software (Microsoft Office Excel®) was employed for statistical analysis. Student’s paired t-tests were also calculated using this package; P-values < 0.05 were considered significant.

**Table 1:** Details of the season, time and outside temperature during the transportation of cartilage biopsy specimens in a TGP cocktail

| **Specimen Ref. no** | **Season of Transport** | **Time of the day** | **Outside Temperature\*** |
| --- | --- | --- | --- |
| 1 | Summer | 1500-1610 hrs JST | High: 27°C; Low: 25°C |
| 2 | Summer | 1400-1500 hrs JST | High: 28°C; Low: 25°C |
| 3 | Summer | 1200-1330 hrs JST | High: 25°C; Low: 24°C |
| 4 | Summer | 1200-1330 hrs JST | High: 22°C; Low: 21°C |
| 5 | Summer | 1300-1400 hrs JST | High: 35°C; Low: 31°C |
| 6 | Autumn | 1430-1530 hrs JST | High: 16°C; Low: 14°C |
| 7 | Winter | 1500-1630 hrs JST | High: 15°C; Low: 13°C |
| 8 | Winter | 1300-1430 hrs JST | High: 10°C; Low: 7°C |
| *Recorded by Weather station at the Tokyo Heliport, Japan (Ref: https://www.timeanddate.com/);<br>JST: Japan Standard Time |  |  |  |

**Table 2:**
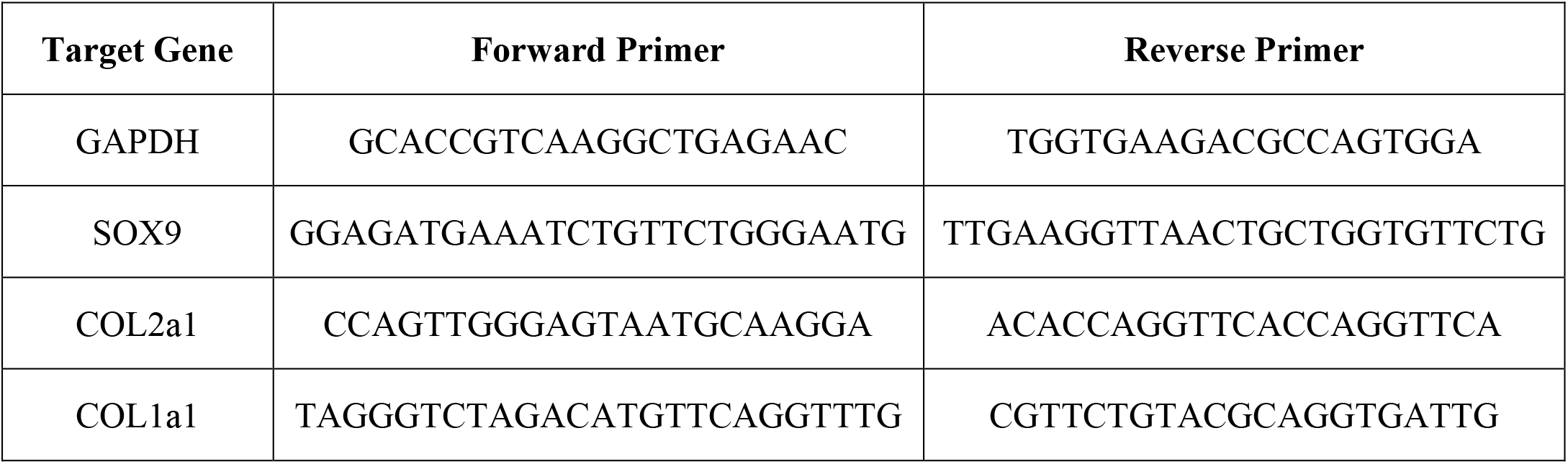
Primer sequence used in RT-PCR

## Results

The weight of the cartilage tissues obtained after biopsy ranged from 0.1 to 0.3 g. The average weight was 0.2 g, which was similar in the ECS- and TGP-transported groups. The cell counts after the isolation of tissues transported in ECS and TGP are given in Figure 2A. The comparison of cell counts obtained after transport of cartilage biopsy tissues correlating to the weight of the cartilage tissue is given in Figure 2B. The cell counts obtained from TGP-transported cartilage biopsy tissues ranged from 0.05 million to 0.4 million cells (average = 0.2 million cells). The cell counts obtained from ECS-transported cartilage biopsy tissues ranged from 0.01 million to 0.3 million cells (average = 0.08 million cells). Thus, the cell counts obtained from TGP-transported tissues were higher than those obtained from ECS-transported tissues. However, this difference was not statistically significant (P = 0.14). The cell counts obtained from TGP-transported cartilage biopsy tissues after 2D monolayer culturing ranged from 0.9 million to 76 million (average = 18 million cells). The cell counts obtained from ECS-transported cartilage biopsy tissues ranged from 0.4 million to 39 million (average = 10 million cells). Thus, the cell counts obtained after 2D monolayer culturing were also higher in TGP-transported tissues compared with ECS-transported tissues. The cells grew as tissues with maintenance of native hyaline phenotype in the 3D-TGP culture (Figure 1E) for a long period of time (six weeks), whereas the cells in 2D monolayer exhibited fibroblast de-differentiation within two weeks (Figure 1F). Cell count could not be performed in 3D-TGP cultures, as the cells grew as tissue-engineered constructs (Figure 1E). RT-PCR results showed that the SOX9 expression was 46.53-fold higher and COL2a1 was 11.6-fold higher in 3D-TGP-cultured tissue than in 2D monolayer cells obtained from tissues transported in TGP. COL1a1 decreased 0.01-fold after 3D-TGP culture compared with 2D monolayer cells obtained from tissues transported in TGP. Expression of SOX9 was 16.22-fold higher and COL2a1 was 10.68-fold higher in 3D-TGP cultured tissue than in 2D monolayer cells obtained from tissues transported in ECS. In addition, COL1a1 decreased 0.01-fold after 3D-TGP culture compared with 2D monolayer cells obtained from tissues transported in ECS (Figure 3).

**Figure 2:**
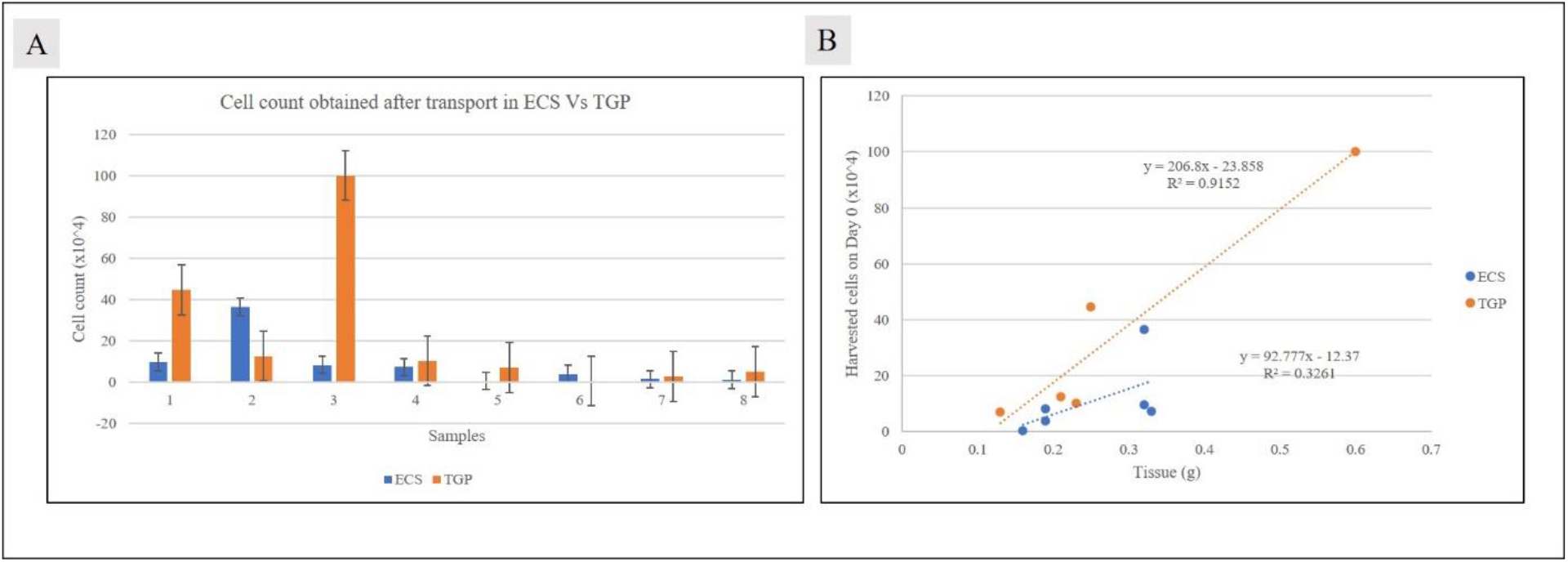
A. Cell count obtained after initial processing of cartilage tissues and after 2D-monolayer culturing for two weeks, B. demonstrating higher cell counts from TGP-transported tissues compared with ECS-transported tissues.

**Figure 3:**
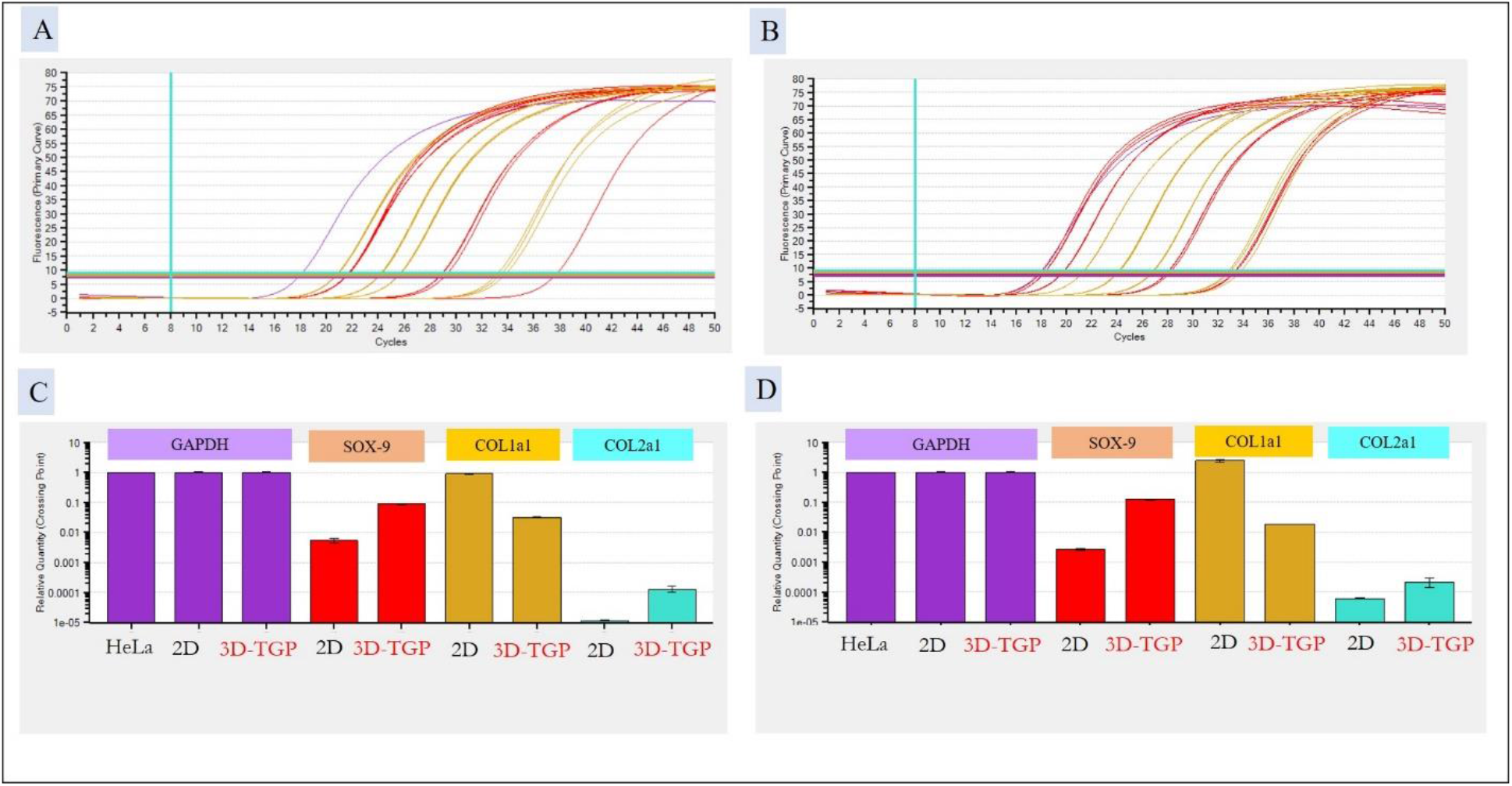
RT-PCR showing expression of SOX9, COL1a1 and COL2a in 2D monolayer-cultured chondrocytes compared to 3D-TGP tissue-engineered cartilage tissue constructs (Figure B and D), with higher expression of SOX9 and Col2a1 higher and lower expression of Cl2a1 in TGP-transported tissues (D) than in ECS-transported tissues (C).

## Discussion

Organ and tissue preservation has its origins in early 20th-century perfusion systems that used simple syringe-based methods in vitro for organs such as the kidney and liver [12]. Later, solutions enabling the transport of organs were developed. One such pioneering effort is that of Collins et al. [13], which is a simple crystalloid with high potassium and low sodium concentrations mimicking the intracellular environment. The solution underwent modifications such as the removal of magnesium, after which it was named Euro-Collins solution (ECS) [14]. Belzer’s team developed another solution [15] with improved buffering and anion composition, which became known as the University of Wisconsin (UW) solution. The ECS and UW solutions remain gold-standard and widely used solutions for organ preservation worldwide [13].

In the case of tissue transport for cell therapies, culture mediums such as DMEM and PBS are widely used. ECS, UW solution, DMEM and PBS all require cool preservation at 4 °C [2]. It should be noted that cold-chain maintenance remains a major hurdle in developing countries and during overseas transport [16]. Although continued efforts are directed at developing potential alternatives, we previously reported the transportation of highly sensitive tissues such as the human corneal endothelial and retinal pigment epithelial tissues without cool preservation over long distances [3,4]. Therapy-applicable cells have been developed after such transport, and these cells have also been transplanted into human patients [5].

According to the Osteoarthritis Research Society International (OARSI), Osteoarthritis (OA) is a highly prevalent debilitating disease affecting 240 million people worldwide. Reports estimate that by 2050, 130 million people will suffer from OA worldwide, and 40 million of these will be severely disabled [17]. Existing approaches to cartilage repair, including abrasion arthroplasty procedures, osteochondral transplantation and ACI, fail to produce repair tissue with the same mechanical and functional properties of native articular cartilage [18]. MACI, which uses scaffolding, has the potential to overcome this hurdle [19]. Both ACI and MACI cell therapies mandate tissue transportation from the source of collection to the cell processing laboratory. TGP has been proven to maintain the native characteristics of cells during transport and culture for a long period of time [3,4,8-10].

This study has clearly established it as an effective technique for maintaining the native hyaline phenotype during transportation, as revealed by the higher cell count in TGP-transported tissues than in ECS-transported tissues (Figure 1) and 3D-TGP-cultured cartilage tissue constructs having higher expression of SOX9 and COL2a1 and lower expression of COL1a1 (Figure 3). It is noteworthy that SOX9 and COL2a1 are necessary for homeostasis in adult cartilage, and their expression is decreased in OA-affected cartilage [20,21]. This higher expression of SOX9 and COL2a1 and lower expression of COL1a1 is more profound in TGP-transported tissues (Figure 3D). It should be noted that the tissues used in the current study were inflamed osteoarthritic tissues. Although re-differentiation of de-differentiated fibroblasts in chondrocyte cultures has been reported when 3D scaffolds are used, such cultures were from normal, non-weight-bearing cartilage. Osteoarthritic cartilage was previously reported to have no significant value for deriving clinically usable chondrocytes [21, 22]. However, the current study, in line with our earlier report [10], has proven that even osteoarthritic cartilage from elderly donors (> 65 years) can be used to generate good-quality cartilage tissue in vitro when the 3D-TGP platform is used, and this outcome is enhanced by using TGP for transportation.

## Conclusion

Transportation of human cartilage tissues in TGP yielded a greater quantity of cells after in vitro processing compared with ECS, and the 3D-TGP-based in vitro culture environment yielded chondrocytes expressing the hyaline cartilage phenotype, as proven by higher SOX9 and COL2a1 and lower COL1a1 expression. Having proven that even OA-affected elderly donor cartilage can be used in this TGP-based transportation and culture methodology to grow good-quality chondrocytes worthy of clinical application to address cartilage damage in cell-based therapies, it is worth recommending a clinical translation of this methodology to improve the outcome of regenerative medicine applications for cartilage repair.

## Declarations

### Ethics approval

The institutional ethics committee of Edogawa Hospital, Tokyo, Japan approved the study. All procedures performed were in accordance with the ethical standards of the institutional ethics committee of Edogawa Hospital, Tokyo, Japan and with the 1964 Helsinki declaration and its later amendments or comparable ethical standards.

### Consent for publication

Informed consent was obtained from all participants and/or their legal guardians whose data has been included in the study.

### Availability of data and materials

Data sharing is not applicable to this article as no datasets were generated or analysed during the current study.

### Authors’ contributions

SK, HY and SA contributed to conception and design of the study. RS helped in data collection and analysis. SA & SP drafted the manuscript. SS, HN and MI performed critical revision of the manuscript. All the authors read, and approved the submitted version.

## Acknowledgements

The authors wish to acknowledge Ms. Takako Fujisaki, Ms. Emi Nagahama & Ms. Junko Tomioka of Edogawa Hospital, Tokyo, Japan for their assistance in sample collection and documentation; Ms Eiko Amemiya and Ms. Sayaka Shimizu of II Dept of Surgery, Yamanashi University, Japan for their assistance with literature collection; Dr. Fumihiro Ijima of Hasumi International Research Foundation, Asagaya, Tokyo, Japan for his assistance with the cell culture work described in the manuscript; Loyola ICAM College of Engineering Technology (LICET) Chennai, India for their support to our research work.

